# Decoding the microstructural properties of white matter using realistic models

**DOI:** 10.1101/2020.06.23.127258

**Authors:** Renaud Hédouin, Riccardo Metere, Kwok-Shing Chan, Christian Licht, Jeroen Mollink, Anne-Marievan Cappellen van Walsum, José P. Marques

**Affiliations:** Radboud University, Donders Institute for Brain, Cognition and Behaviour, Nijmegen, Netherlands; Radboud University Medical Centre, Medical Imaging and Anatomy, Nijmegen, Netherlands; Computer Assisted Clinical Medicine, Medical Faculty Mannheim, Heidelberg University, Germany; Empenn, INRIA, INSERM, CNRS, Université de Rennes 1, Rennes, France

**Keywords:** White matter models, Microstructural properties, Magnetic susceptibility, Deep learning network

## Abstract

Multi-echo gradient echo (ME-GRE) magnetic resonance signal evolution in white matter has a strong dependence on the orientation of myelinated axons with respect to the main static field. Although analytical solutions have been able to predict some of the white matter (WM) signal behaviour of the hollow cylinder model, it has been shown that realistic models of WM offer a better description of the signal behaviour observed.

In this work, we present a pipeline to (i) generate realistic 2D WM models with their microstructure based on real axon morphology with adjustable fiber volume fraction (FVF) and g-ratio. We (ii) simulate their interaction with the static magnetic field to be able to simulate their MR signal. For the first time, we (iii) demonstrate that realistic 2D WM models can be used to simulate a MR signal that provides a good approximation of the signal obtained from a real 3D WM model derived from electron microscopy. We then (iv) demonstrate *in silico* that 2D WM models can be used to predict microstructural parameters in a robust way if ME-GRE multi-orientation data is available and the main fiber orientation in each pixel is known using DTI. A deep learning network was trained and characterized in its ability to recover the desired microstructural parameters such as FVF, g-ratio, free and bound water transverse relaxation and magnetic susceptibility. Finally, the network was trained to recover these micro-structural parameters from an *ex vivo* dataset acquired in 9 orientations with respect to the magnetic field and 12 echo times. We demonstrate that this is an overdetermined problem and that as few as 3 orientations can already provide comparable results for some of the decoded metrics.

[Highlights] - A pipeline to generate realistic white models of arbitrary fiber volume fraction and g-ratio is presented; - We present a methodology to simulated the gradient echo signal from segmented 2D and 3D models of white matter, which takes into account the interaction of the static magnetic field with the anisotropic susceptibility of the myelin phospholipids; - Deep Learning Networks can be used to decode microstructural white matter parameters from the signal of multi-echo multi-orientation data;

## 1. Introduction

White matter (WM) consists mainly of myelinated axons and plays an important role in the transmission of information across the brain. The myelin sheath surrounding axons acts as an electrical insulator, thus increasing the transmission speed of the nerve impulses. The development of myelin played a key role in evolution and the emergence of large vertebrates [1] and it is still central to brain maturation [2]. The degradation of myelin, commonly referred to as demyelination, is present in various neurodegenerative diseases and can lead to severe motor and mental disabilities [3]. Such neurodegenerative disorders (e.g multiple sclerosis) show high variability among individuals, and it is difficult to predict and understand the course of the disease by solely counting the number of lesions or comparing the values obtained in magnetic resonance (MR) relaxometry [4]. Therefore, non-invasive imaging methods that can investigate the WM microstructure such as myelination may offer important means to study neurodegenerative diseases, providing crucial information for diagnosis, and monitoring progress and assessment of potential treatment effectiveness.

Direct MR imaging of the myelin is challenging due to the ultra-short transverse relaxation time of the phospholipid proton 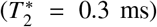 [5]. Nevertheless, several attempts have been performed using zero or ultra-short echo time techniques [5, 6]. Myelin can be probed indirectly using magnetization transfer techniques [7, 8]. Alternatively, myelin water imaging is a method that attempts to measure the signal of water that is trapped in between myelin layers and that was originally based on multi-echo spin-echo data [9] and has more recently been explored using multi-echo gradient-echo data [10]. However, the detection of myelin water remains challenging due to its short *T*_2_ value (∼20 ms) and 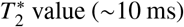 [11]. In this paper, we will focus on myelin water imaging using a multi-echo gradient echo (ME-GRE) sequence.

WM is a complex environment composed of not only axons but also different types of glial cells, vessels and more. However, the biophysical models typically used in magnetic resonance imaging (MRI) are reduced to 3 compartments: intraaxonal, myelin and extra-axonal water protons. Axons in WM have various shapes and sizes, with a diameter ranging from 0.1 *µ*m to 2 *µ*m for unmyelinated axons and from 1 *µ*m up to 10 *µ*m for myelinated axons [12], and are typically modelled as cylinders. The myelin sheath, formed in the central nervous system (CNS) by oligodendrocytes, represents approximately 80% of the brain’s dry weight and consists of tightly packed phospholipid bi-layers united by the hydrophobic tails, separated by water layers [13]. These phospholipids, because of their elongated form and their radial organisation around the axon, have an anisotropic magnetic susceptibility [14, 15] with diamagnetic property when compared to the surrounding water. These microstructural features are believed to be well approximated by a tensor with cylindrical symmetry which can be expressed as a sum of an isotropic (*χ*_*i*_) and anisotropic (*χ*_*a*_) components. Various values have been reported in the literature of myelin for *χ*_*i*_ ranging from 0.13 to 0.06 ppm and *χ*_*a*_ ranging from 0.15 to 0.09 ppm [16, 17, 18] (with ppm considered with respect to the magnetic susceptibility of pure water).

In the presence of a strong magnetic field, a secondary microscopic magnetic field perturbation is created by these phospholipids [19]. This secondary field can be observed in both magnitude and phase of a ME-GRE signal [20]. One manifestation of the anisotropic magnetic susceptibility of myelin is that the MR signal of a GRE sequence shows a dependence on the orientation of the fibers relative to the main magnetic field. It has been shown that simple 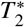 maps are orientation dependent [21], and hence unsuitable for the estimation of myelin properties. Part of this orientation dependence can be accounted for using *a priori* knowledge of fiber orientations [22].

In this study, we set out to investigate the feasibility of WM microstructure property quantification using realistic WM fiber geometries, which has the potential to measure microstructure properties without the bias associated with simplification of the biological environment in analytical models. Firstly, we developed a method to generate hypothetical 2D WM models based on realistic axon shapes. ME-GRE signals with different axon and myelin properties were subsequently simulated using these 2D WM models. The validity of these 2D models was tested by comparing the signal similarity between the signal simulated from a 3D WM model (obtained by 3D electron microscopy of a genu of a sagittal mouse corpus callosum section) to that of 2D models with matched microstructural parameters. Secondly, a dictionary of ME-GRE was simulated using realistic WM models with a wide range of WM microstructure properties. This dictionary was then used to train a deep neural network to recover WM microstructure properties from ME-GRE signal. ME-GRE signal with multiple object orientations with respect to the main magnetic field data was used in this process to ensure there is sufficient signal variation due to the susceptibility properties of myelin. Finally, we validated and optimised this deep neural network using *in silico* data and applied the same method on *ex vivo* data. This process is briefly summarized in Fig. 1.

**Figure 1:**
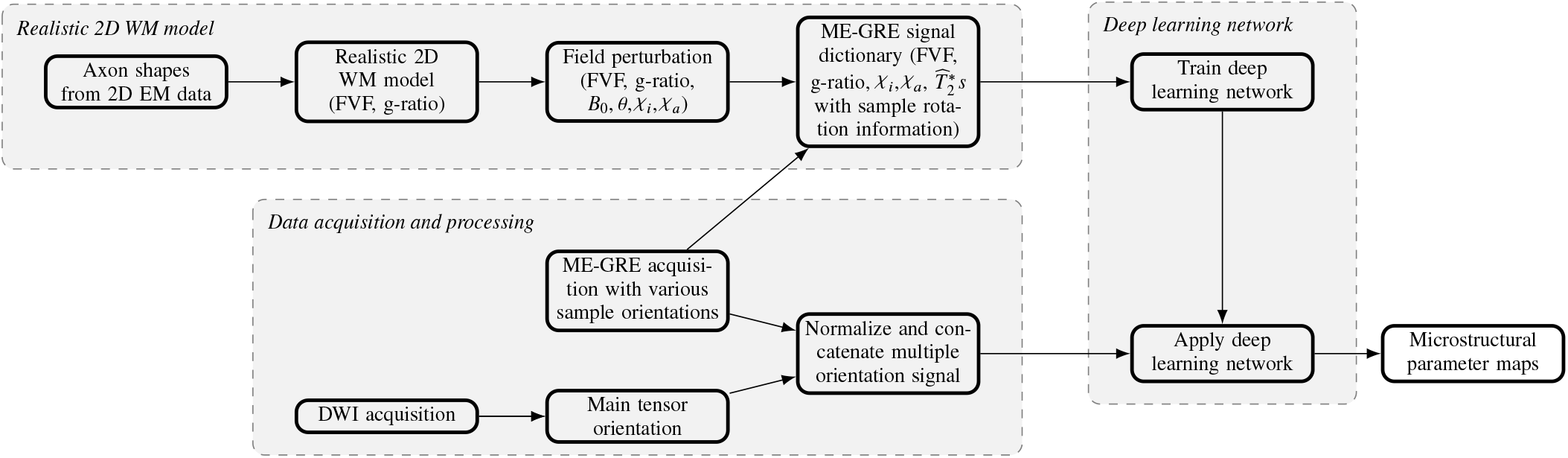
Diagram of the entire pipeline from the data acquisition and the creation of 2D realistic WM models to the recovery of microstructure parameter maps.

## 2. Methods

### 2.1. 2D WM model

In the presence of magnetic field, the magnetic susceptibility of myelin relative to its surrounding creates a secondary magnetic field, which although small, affects the MRI signal both in phase and magnitude. These phenomena have been used in the past to study WM orientation [23, 22] and can be studied both analytically and numerically considering various simplified WM models.

#### 2.1.1. Hollow cylinder model (HCM)

The HCM, proposed by Wharton and Bowtell, is commonly used to approximate WM microstructure [17]. The myelin sheath is represented by an infinite hollow cylinder with an inner radius *r*_*i*_ and an outer radius *r*_*o*_. The inner part of the hollow cylinder is the intra-axonal compartment and the external part is referred as the extra-axonal compartment.

This 3-compartment cylindrical representation of WM allows an analytical derivation of the field perturbation in each of those regions and characterization WM using:

- Fiber volume fraction (FVF) - the proportion of myelinated axon within the model
- g-ratio - the ratio between the intra-axonal radius (*r*_*i*_) and the myelinated axon radius (*r*_0_):

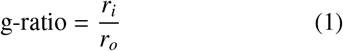

This solution, which is very convenient to model, offers, for example, an analytical estimation of the fiber-orientation dependence of 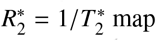 map[23].

However, it has been recently demonstrated that the HCM has intrinsic biases compared to a more realistic WM model created from electron microscopy data [14]. The circular axon shapes create artificially large frequency peaks, in particular within the intra-axonal compartment, which are not present in a realistic model. In the following section we will present the creation of a realistic 2D WM model based on real axon shapes and realistic size distributions.

#### 2.1.2. Electron microscopy based models

In this study, we used a 2D electron microscopy image of an entire slice of a canine spinal cord from an histology open database ^1^ [24] as our database of axon shapes. The sample is 5mm width and 7.5mm long with a 0.25*µ*m resolution which corresponds to a 20.000 × 30.000 image. An open-source segmentation software was used to segment the image leading to a collection of ∼600.000 myelinated axon shapes [25]. The resulting axons had an average diameters of 2.9 ± 0.1 pixels and g-ratio of 0.62 ± 0.01. The resolution is sufficient because we do not want to segment unmyelinated axons that have been shown to have no significant impact on the ME-GRE signal obtained [14]. The unmyelinated axons are therefore included within the extra-axonal space. In case of a realistic axon shape, the g-ratio is redefined as the square root of the ratio between the intra-axonal surface and the outer surface (measured as the number of myelinated pixels with at least one side in direct contact with intra or extra-axonal space).

#### 2.1.3. Axon packing algorithm

A set of 400 axon shapes was randomly picked from the the collection above to create a realistic 2D WM model with predefined FVF and g-ratio. To do so, we developed an axon packing algorithm based on an existing software [26] that had been initially developed for cylindrical axon models. The packing process is performed as follow (see Fig. 2):

**Figure 2:**
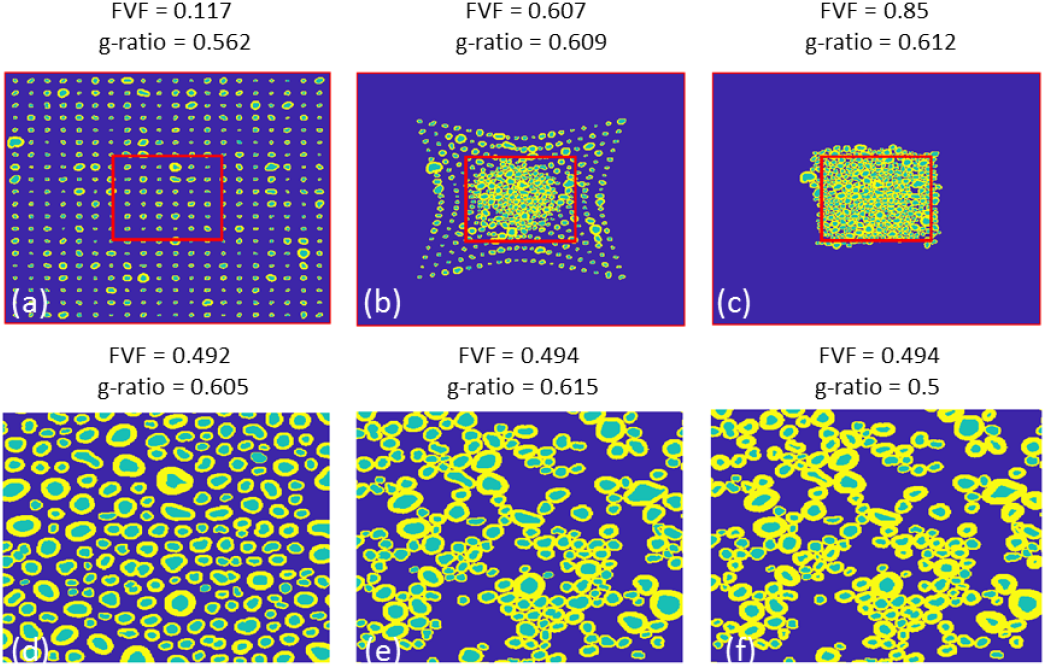
Top row: 400 axons are placed on a grid (a) and packed following an attraction/repulsion method (b) until high FVF is reached (c). Bottom row: Zoom on the mask delineated by the red square. A desired FVF is reached by spreading the axons from the center (d) or randomly removing some axons (e). Keeping the same axons and thus the same FVF, the myelin thickness can be modified to obtain an expected g-ratio (f)

##### Algorithm 1: Axon packing

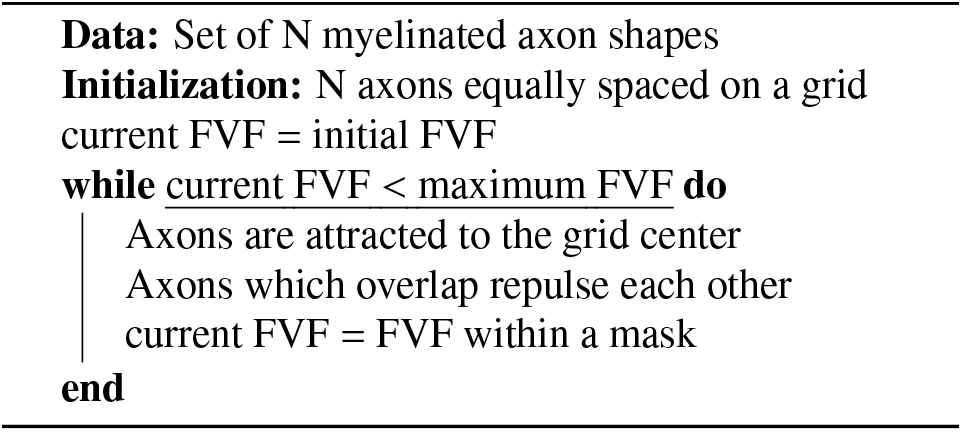

In the current implementation, as the axon shapes are picked randomly, they do not necessarily fit optimally together (during the attraction and repulsion process, the axon is not allowed to rotate), thereby generating small gaps within the model. The maximum FVF parameter, corresponding to a model where the axons are highly packed while avoiding overlap was empirically found to be 0.85. According to literature, such an FVF value already represents a WM model with a very high axon density [27].

#### 2.1.4. Obtaining an expected FVF

Once the maximum FVF for a given collection of axons is achieved, this packed WM model was used to obtain a new model with an a different FVF. Two different methods, illustrated in Fig. 2, were proposed: (i) randomly remove axons or (ii) spread the axons from the figure center. The first method creates important gaps within the extra-axonal space that could correspond to glial cells or bundles of unmyelinated axons, while the second method creates a more uniformly distributed WM model. Based on the EM data visually explored up to now, both could be valid representations. Their corresponding field perturbation histograms were close enough and both models were used to enforce the diversity of our WM model dictionaries.

#### 2.1.5. Change the g-ratio

Finally, the mean g-ratio of the model was modified, while keeping the FVF constant. This operation was performed on an axon-by-axon basis by dilating or eroding the inner myelin sheath by one pixel, to ensure a smooth modification of the g-ratio, depending on whether the g-ratio was to be decreased or increased. Each axon has a given probability to be randomly picked, this probability is linked to its diameter. As the dilatation/erosion is fixed to one pixel, larger axons need to be picked more frequently to respect the original proportion of FVF. It should be noted that one axon can be selected multiple times for erosion. The modification of the g-ratio is illustrated in Fig. 2 and a video of the entire 2D WM model creation is available as supplementary material, where it can be seen that a given axon can be selected multiple times. Eventually, different models with similar FVF and g-ratio can be created using our large axon shapes database and the code made available in the toolbox.

### 2.2. Signal creation

With a view to using these 2D models to simulate the ME-GRE signal, we need to define the susceptibility of pixel element, compute the induced magnetic field perturbation and eventually simulate the signal evolution in this inhomogeneous environment.

#### 2.2.1. Magnetic susceptibilities

For the sake of simplicity, we consider the intra-axonal and extra-axonal compartments to have equal magnetic susceptibility, for it to be isotropic and to have value zero. As a result, the susceptibility attributed to myelin is the difference between the myelin susceptibility and the susceptibility of the surrounding compartments. In the myelin compartment the magnetic susceptibility is described by a tensor that results from the sum of an isotropic (*X*_*i*_) and an anisotropic (*X*_*a*_) component:

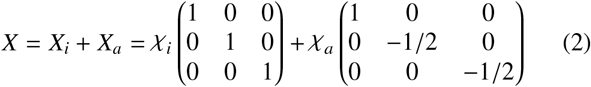

where *χ*_*i*_and *χ*_*a*_are scalar isotropic and anisotropic susceptibility multiplicative constants, respectively. The susceptibility tensor *X*_*R*_ within the myelin sheath in a 2D model is determined by the phospholipid orientations *ϕ* with respect to the magnetic field on that plane [17]:

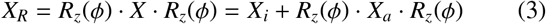

with *R*_*z*_ (*ϕ*) the 3D rotation matrix around the z axis.

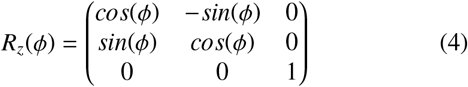

it should be noted that the tensor map only depends on the phospholipid orientation (see Fig. 3) and is not related to the main magnetic field. In simple cases, as for the HCM, the computation of *ϕ* is trivial. For more complex axon shapes, there is no straightforward definition of the orientation of the phospholipids throughout the whole myelin sheath. Its orientation should be perpendicular to the tangent of the myelin surface, but the inner and the external boundaries are neither smooth nor necessarily parallel to each other in our segmented models. The orientation of the phospholipids is estimated on an axon-by-axon basis. First, the selected axon is placed in a small matrix (including 10 pixels of each side of the axon edges for computational time considerations), then the extra-axonal, myelin and intra-axonal compartments are given the values of 0, 1 and 2 respectively. The resulting map is smoothed with a 2D Gaussian filter with a width of 5 × 5 to create a smoothed pyramidal structure. If the myelin sheath is too large and still contains piecewise constant part after smoothing, the process is repeated. Finally a 2D gradient is computed from the smoothed map. As the map is smoothly varying from 0 to 2 within the myelin compartment, the gradient at each point will define the steepest direction from the extra- to the intra-axonal space, the phospholipid orientation is assumed to correspond to the gradient direction (see Fig. 3b).

**Figure 3:**
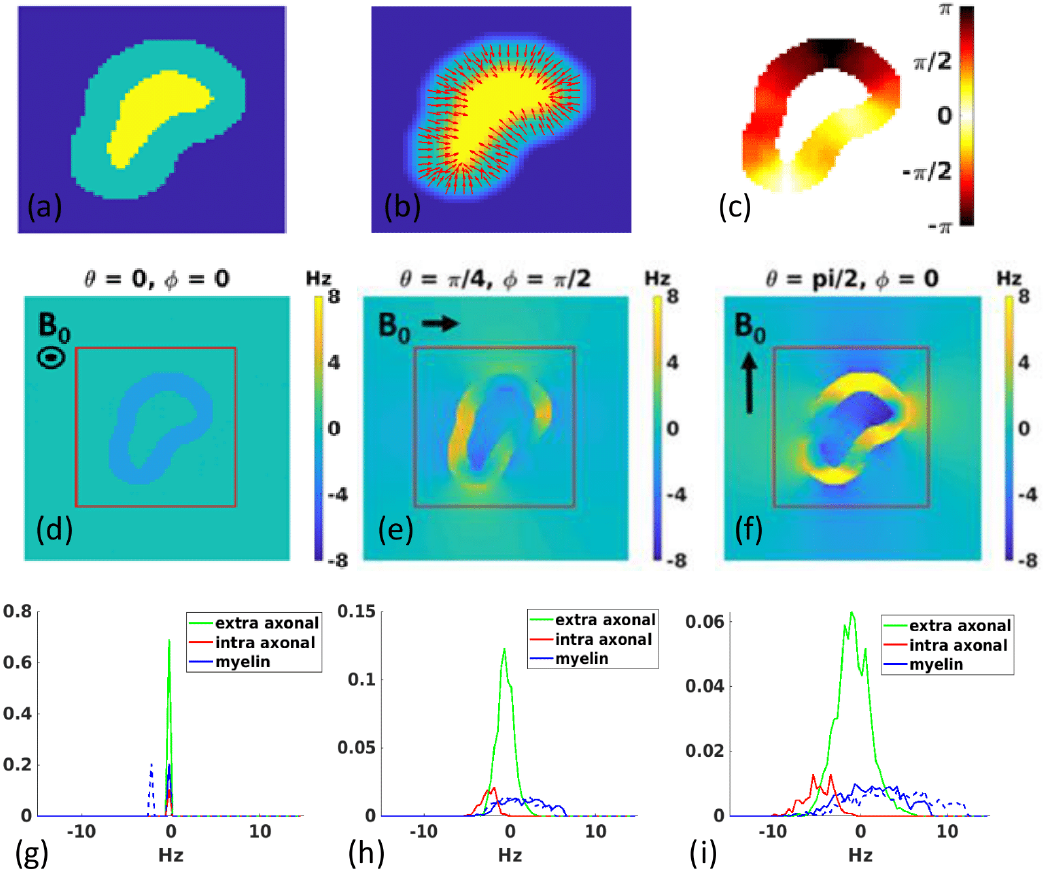
First row represents the phospholipid orientation estimation performed for each axon. (a): the extra-axonal, myelin and intra-axonal compartments being assigned values 0, 1 and 2 respectively. (b): the model is smoothed with a Gaussian filter (c) Gradient orientation is computed on the smoothed map. Second row: Field perturbation for one axon with 3 different magnetic field orientations. Third row: Corresponding histograms computed within the red square to keep a reasonable FVF. The myelin histogram is presented with Lorentzian correction (solid blue line) and without Lorentzian correction (dashed blue line). A magnetic field parallel to the axon orientation can be characterized by Dirac delta functions with a value of 0 for the intra and extra axonal compartments and a negative or null value for the myelin compartment. A perpendicular magnetic field creates much stronger perturbations and present broad distributions within the 3 compartments.

#### 2.2.2. Field perturbation

From the phospholipid orientation map, the susceptibility tensor map can be calculated using Eq. 3. The susceptibility tensor map is used to compute the field perturbation in the frequency domain as described in [28]. An illustration of the field perturbation generated by a single axon for several *B*_0_ orientations, with and without the Lorentzian correction (see Section 2.2.3), is shown in Fig. 3. The induced field perturbation strongly depends on the *B*_0_ orientation. A magnetic field parallel to the axon orientation has a small negative field shift or no field shift at all within the myelin sheath while a perpendicular magnetic field creates much stronger perturbations within the 3 compartments. The overlapping frequency spectra of the 3 compartments make them hard to disentangle.

#### 2.2.3. ME-GRE signal

In previous studies the ME-GRE signals was computed as [16]:

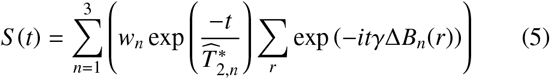

where 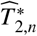 is the compartment specific apparent transverse relaxation rate that is not originated by myelin induced field inhomogeneities and *w*_*n*_ is the water weight (reflecting the water signal per pixel in our 2D model, which includes proton density and T1 saturation effects). The field perturbation Δ*B*_*n*_(*r*) at each pixel (computed considering the sphere of Lorentz assumption) is therefore responsible for the signal decay associated with myelin induced field inhomogeneities, 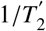, contributing to the each compartments’ apparent transverse relaxation rate 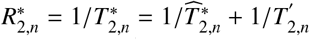.

In our implementation a correction has been introduced in the frequency shift of the myelin water compartment to account for the compartmentalization of water. Instead of using the standard Lorentzian sphere approximation used for the field computation, we have used the cylindrical Lorentzian approximation [29] similar to the initially proposed by He and Yablonskiy. This correction was done separately for each pixel within the myelin compartments and taking into account the susceptibility tensor, such that:

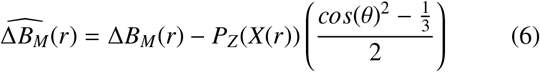

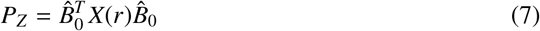

where 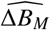 is the Lorentzian corrected myelin field perturbation and *P*_*Z*_ is the projection of the susceptibility tensor along the *B*_0_ orientation. An illustration of the ME-GRE signals simulated with the Lorentzian correction is shown with two examples of WM geometry in Fig. 4. Intuitively, in this formalism, the myelin sheath is broken into various infinite cylinders running parallel to the axon. For a closer inspection to the impact of this correction on the frequency of the myelin water compartment as a function of axon orientation for the more tractable case of a cylinder, refer to Appendix A. There, we also compare the current correction to the more advanced layered models [15, 30] and discuss the pros and cons of the different approaches.

**Figure 4:**
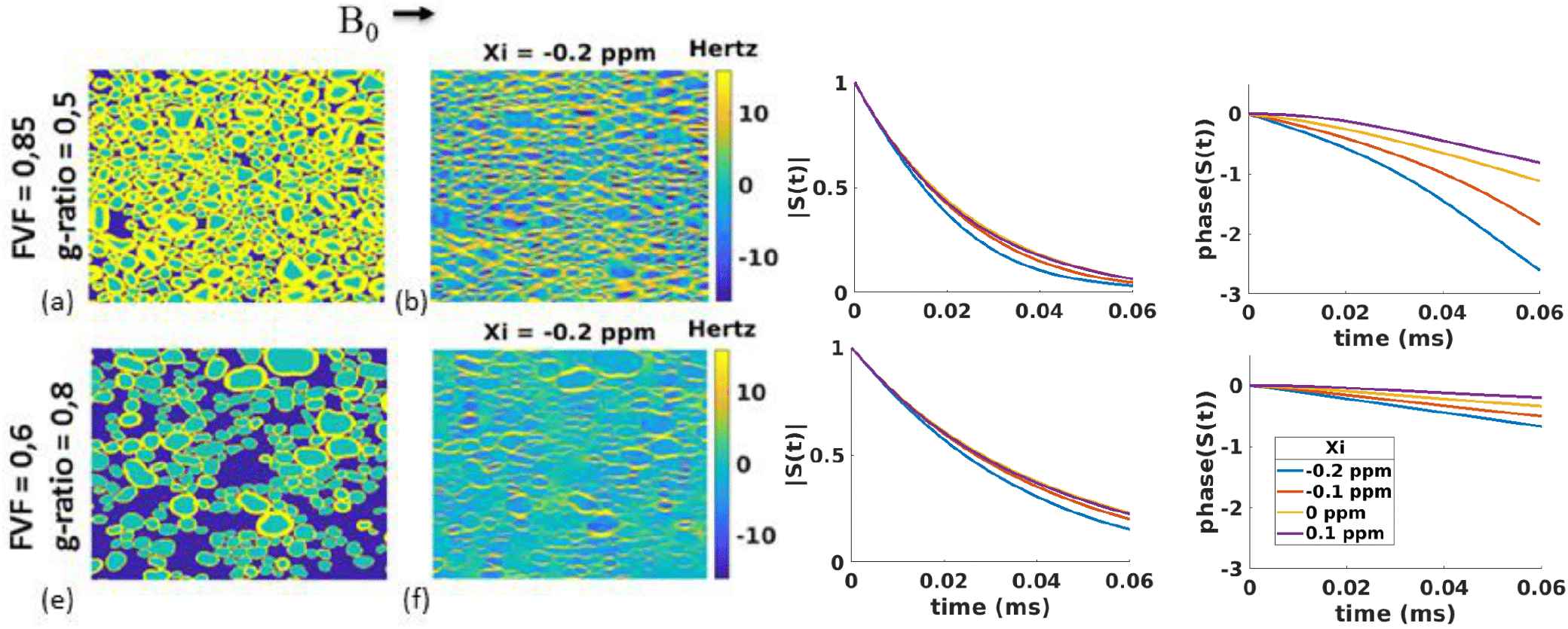
Variations of signal decay as function of FVF, g-ratio and susceptibility: (a,e) Two examples of realistic WM models with different FVF and g-ratio (b, f) corresponding field perturbation when axons perpendicular to the magnetic field; Simulated ME-GRE signal with the Lorentzian correction (c, g) magnitude and phase (d, e) for the two models with different isotropic susceptibility. Remaining model and relaxation parameters are fixed according to literature values (see Table 1).

MRI data amplitude depends not only on the magnetization amplitude, but also on the RF coil sensitivity and receiver gain. The phase depends on the RF transceiver and on the quality of the B0 shimming and presence of fields due to the susceptibility of neighbouring pixels. To be able to compare our simulations to real data, both the simulated and measured signals were normalized as follows :

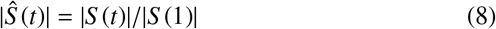

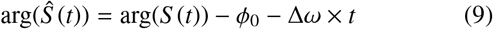

where arg(*Ŝ* (*t*)) is the phase of the signal, *ϕ*_0_ and Δ*ω* are the phase and frequency offsets estimated by performing a simple linear regression on arg(*Ŝ*(*t*)). Note that it is relevant to also perform such a normalisation in the simulated data as its amplitude and frequency would depend on the myelin volume fraction. At a later stage, when training a network to decode microstructural parameters, it is important to ensure the network is trained on signal features that are experimentally relevant, such as non-exponential behaviour of the signal decay and non-linear phase evolution.

#### 2.2.4. Model validation

While the realistic 2D WM models have been shown to better represent the ME-GRE signal of WM than the simple HCM, they assume the replication of the same structure along the third dimension resulting in bundles that are unrealistically aligned and cannot represent the natural dispersion present in a real axon bundle. Dispersion can occur not only in regions of fiber crossing, fiber kissing, but also in regions traditionally expected to be unidirectional such as the corpus callosum [33]. However, 3D models are hard to construct, not only because of the lack of 3D EM data (that could represent a ground truth), but also because of the complexity of 3D axon packing [34]. Also, the 2D axon shapes used in our realistic WM modeling can possibly be elongated as they are obtained from cutting through axons that were not perpendicular to the surface. Furthermore, the estimation of the susceptibility tensor map and the field perturbation in 3D models would make the process even more time consuming. We have designed a small study, presented in the Appendix B to assess the ability of our 2D models to represent a real 3D model with comparable microstructural properties.

### 2.3. Dictionary creation

A dictionary of signal evolution can be created using the simulated ME-GRE signals in the presence of different WM models. Such dictionary can be used to derive the microstructural tissue properties from the ME-GRE signal by using root-mean-square minimization between the dictionary elements and measured signal, as previously done in, for example, finger-printing [35]. Alternatively, a deep learning network can be trained to learn the tissue properties from the dictionary as will be demonstrated later.

The WM model and the magnetic field distributions present on each of its compartments depend on 5 microstructure related parameters: FVF, g-ratio, *χ*_*i*_, *χ*_*a*_, as well as the fiber orientation. For the purpose of training a deep learning network, we considered repeating simulations with various axon packing using the aforementioned properties. The ME-GRE signal from each WM model depends on the specific NMR properties of each compartment 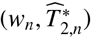. This would result in 6 supplementary parameters. To reduce the dictionary size, the 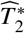 and the water density from the intra- and extra-axonal pixels were defined to be the same. This reduced the number of parameters from 6 to 3: 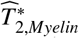 the 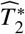 of myelin water; 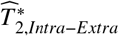 the 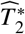 of the intra and extra-axonal compartment and a relative water weight given by *w* = *S* _0,*Intra*−*Extra*_*/S* _0,*Myelin*_. The parameter ranges used to construct the dictionary are presented in Table 1 along with typical WM values. The dictionary has 8 dimensions, with 5 to 20 entries per dimension leading to 7.680.000 vectors. In the following *in silico* and *ex vivo* experiments, all the dictionaries have the same parameter ranges.

**Table 1:** Table describing our 8 dimension dictionary of signal models. First and third column describe the parameters varied and their expected mean values as found in literature. Middle column: Parameter range used in our dictionary, minimum : step : maximum. *The relative water weight in the axon and in the intra and extra-axonal space depends in our case on the acquisition parameters, flip angle and TR. The value presented was the one used for the typical WM deep learning experiment.

| Model parameters | Dictionary | Typical WM values |
| --- | --- | --- |
| FVF | 0.1:0.1:0.8 | 0.7<sup>a</sup> |
| g-ratio | 0.5:0.05:0.85 | 0.65<sup>b</sup> |
| $\chi_i$ (ppm) | -0.2:0.1:0.2 | -0.1<sup>c</sup> |
| $\chi_a$ (ppm) | -0.1 (fixed) | -0.1<sup>a</sup> |
| $\widehat{T}_{2,Intra-Extra}^*$ (ms) | 20:20:100 | 60<sup>d</sup> |
| $\widehat{T}_{2,Myelin}^*$ (ms) | 4:4:20 | 16<sup>d</sup> |
| $w_{Myelin}/w_{Intra-Extra}$ | 0.5:0.5:3 | 2* |
| Number of fiber orientations | 20 | / |

<sup>a</sup> (Choy et al., 2020) [31]
<sup>b</sup> (Mohammadi et al., 2015) [32]
<sup>c</sup> (Wharton et al., 2012) [17]
<sup>d</sup> (Xu et al., 2018) [14]

Each entry of the dictionary is composed of the normalized signal magnitude and phase (or real and imaginary components, 2 x nTE with nTE the number of echo times in the simulation) and an additional entry encoding the fiber orientation information characterized by the angle between the fiber and the static magnetic field. When deriving the microstructural properties from measurements with multiple orientations with respect to the magnetic field, the signal is concatenated along the *n* orientations which leads to a vector size of *n* · (2TE + 1). An illustration of such simulated normalized signals magnitude and phase with different orientations is presented in Fig. 5. Unlike a single orientation dictionary, this multi-orientation dictionary is only valid for a specific set of rotations used in a specific acquisition.

**Figure 5:**
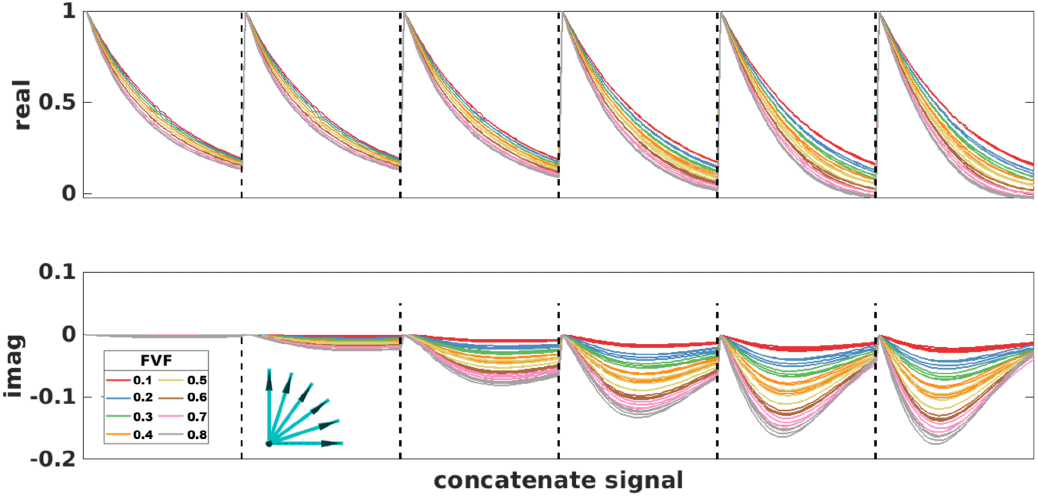
The ME-GRE signal with the Lorentzian correction is simulated with 6 magnetic field orientations, separated by dashed line on the figure, *θ* = 0 −*π/*2 equally space (see arrows), for WM models with different FVF from 0.1 to 0.8 (4 models for each FVF). The top and bottom rows represent respectively the signals real and imaginary part for each of the 6 orientations separated by a vertical black line.

### 2.4. Deep Learning

The ME-GRE signal dictionary was used to train a deep learning network using Keras with TensorFlow GPU backend [36]. For all the following experiments, the dictionaries were trained on 7 entire sets of WM models and assessed by the loss function on another set of WM models, which correspond to a validation split of 0.125. This network is composed with 3 hidden layers of size 2 \**l*_*i*_ \**l*_*o*_, 1.5* *l*_*i*_ \**l*_*o*_, 1.25* *l*_*i*_ \**l*_*o*_, *l*_*i*_ and *l*_*o*_ being the concatenate signal length and the number of parameters, with a respective dropout of 0.4, 0.2, 0.1 using a tanh activation function and an additional linear layer, see Fig. 6. Both inputs and outputs were normalized, a stochastic gradient descent optimizer was used and the loss function is a mean absolute error.

**Figure 6:**
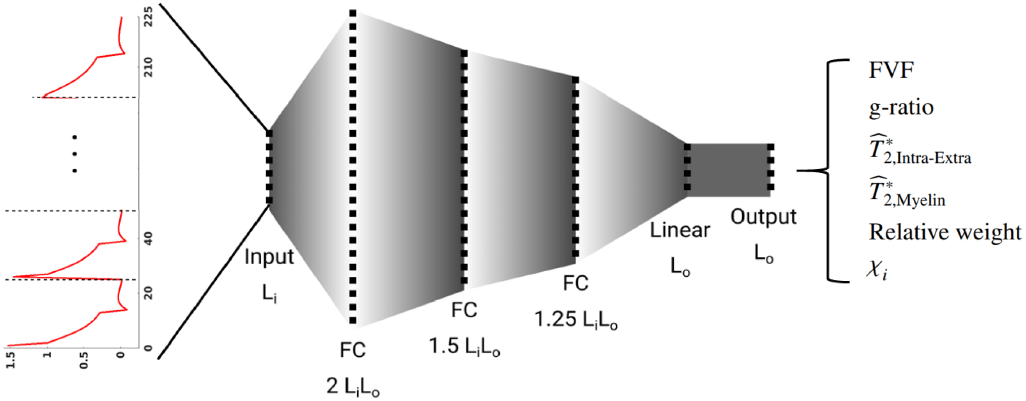
Illustration of the architecture of the deep learning network used in this manuscript. The input is the measured signal (real and imaginary values) together with the main fiber orientation obtained from DTI. The network has 3 hidden layers. FC stands for fully connected layer, Li and Lo are respectively 225 and 6. The output of the network is a vector containing the 6 microstructure parameters. The inputs signals have a vector size of 225 which correspond to the concatenation of 9 orientations. Each orientation includes the *θ* angle between the fiber and the magnetic field orientation, the normalized signal real and imaginary part along 12 TE.

To gain experience on our network ability and limitations to derive microstructure properties, its performance was first tested on numerical simulations. Particularly we wanted to assess what the optimum echo time range and the number of echoes were, as well as study the gains associated with different numbers of sample rotations needed to successfully recover WM properties (which will affect our data acquisition protocol). The design and training of the network were also subjects of careful attention. The deep learning hyperparameters were tuned following an empirical approach, with the selected ones giving results that are both accurate and robust to the change of signal parameters.

The validation loss function (mean absolute error of the parameters estimated on a validation data set - one set of WM models which is not used for training) was used as a metric to assess the convergence of the network. All the parameters, within their range, were re-scaled between 0 and 1, to make validation loss a less arbitrary number. This metric is an average of the mean absolute error for each parameter, thus, it does not allow performing fine comparisons. Despite this remark, the validation loss is a classic and robust way to assess the training process with an unique number.

#### 2.4.1. Deep Learning performance evaluation on simulated data

The robustness of the parameter recovery was tested by adding a complex white noise (0%, 0.5%, 1%, 2% and 4%) to a ME-GRE signal on a dictionary used in the training and validation processes. The noise levels mentioned above are relative to the signal amplitude at the first echo, TE = 2.15 ms. The first 3 columns of Table 2 summarize the parameters used in the creation of the dictionary and training of the network. The rotations used were chosen to mimic the experimental protocol used on an *ex vivo* acquisition described later in this section.

**Table 2:**
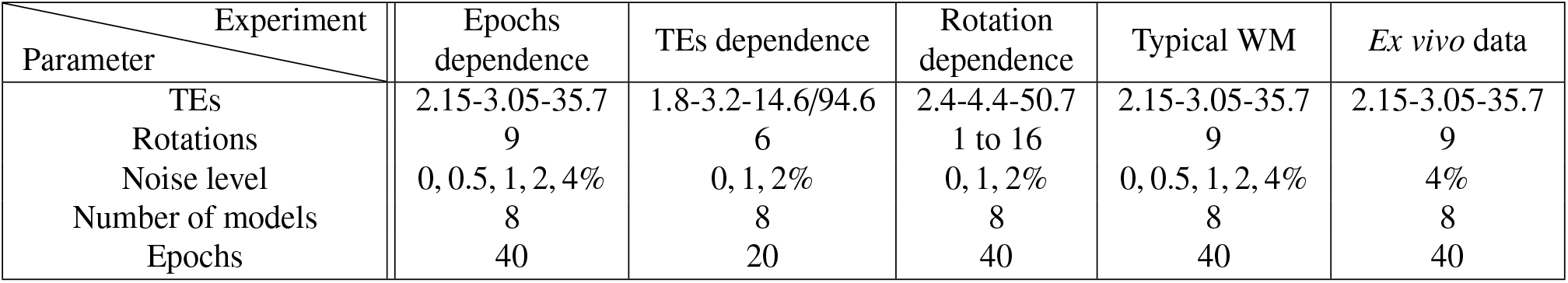
Table describing the parameters used in the training of each of the experiments described in the Methods section. This include parameters associated with the dictionary (echo times, number of rotations, noise level, number of independent WM models) and a deep learning parameters (number of epochs). The four first columns refer to the *in silico* experiments and the last column corresponds to the real *ex vivo* experiment. It should be noted that in the case of the entries with 9 orientations, these were same 9 orientations and were derived from the rotations obtained from the co-registration of the ex-vivo sample

| Parameter \ Experiment | Epochs dependence | TEs dependence | Rotation dependence | Typical WM | *Ex vivo* data |
| --- | --- | --- | --- | --- | --- |
| TEs | 2.15-3.05-35.7 | 1.8-3.2-14.6/94.6 | 2.4-4.4-50.7 | 2.15-3.05-35.7 | 2.15-3.05-35.7 |
| Rotations | 9 | 6 | 1 to 16 | 9 | 9 |
| Noise level | 0, 0.5, 1, 2, 4% | 0, 1, 2% | 0, 1, 2% | 0, 0.5, 1, 2, 4% | 4% |
| Number of models | 8 | 8 | 8 | 8 | 8 |
| Epochs | 40 | 20 | 40 | 40 | 40 |

The ME-GRE signal of a given WM model depends on the magnetic field orientation with respect to its structure (see Fig. 5), this leads us to adopt a multi-orientations approach when trying to decode WM microstructure properties. However, as an increased number of orientations means a longer acquisition time, we performed a theoretical comparison study to estimate the benefit of using a large number of orientations vs a reduced number of orientations with data that has higher SNR. A dictionary with 16 optimal orientations was created for 3 different noise levels (0, 1 and 2%). In order to maximize information, each fiber should have the largest possible range of *θ* from 0 to *π/*2. To do so, the 16 3D rotations had evenly spread axis on the sphere with a common *π/*2 angle. Then, for a range of number of orientations from 1 to 16, a subset of this dictionary was used to train a deep learning network.

The influence of the number of echoes on the deep learning parameter recovery performance was tested. To do so, several networks were trained with a fixed echo spacing (3.05 ms - mimicking our experimental protocol), a various number of TEs (5, 10, 15, 20, 25 and 30) and noise levels. At this stage no considerations of the impact on *T*_1_ weighting were factored into the analysis.

Finally, we tested the deep learning for one set of realistic parameter values of WM (see Table 2), that allows to detail the behavior of each parameter individually. The signal was simulated 125 times for 8 independent WM models leading to 1000 signal simulations with each different noise level. We tested two methods to recover the parameters: (i) using a deep learning network trained with a noise matching the simulated noise; (ii) using a deep learning trained with a maximum noise level regardless of the simulated signal noise.

### 2.5. Ex vivo data acquisition

A formalin fixed post-mortem brain (female, 88 years old, 26 hours of post-mortem interval and 7-month fixation period) was scanned in a 3T scanner (Prismafit, Siemens, Germany).490 The brain was scanned in 9 orientations relative to the static magnetic field. To avoid brain deformation between different rotations, a customised 3D brain holder was built and used throughout the scanning session [37]. Prior to scanning, formalin was washed out using distilled water and prepared in low pressure environment, using a vacuum pump at 20 mbar during 12h to remove all air bubbles trapped in the various cortical sulci. During this period the brain was occasionally rotated to ensure removal of air trapped inside the ventricles.

For each head position the following protocol was repeated:- (a) 3D monopolar ME-GRE with 12 echos (TE = 1.7 : 3.05 : 35.25ms, TR = 38 ms), with a 1.8mm isotropic resolution and matrix size (128×128×128), acquisition time 8.21 mins. This protocol was repeated 6 times with 6 different flip angles (*α* = 5° */* 10° */* 15° */* 20° */* 35° */* 65°); -(b) an MP2RAGE [38] with 1mm isotropic resolution was acquired for co-registration purposes. The MP2RAGE parameters were adapted to be able to map the short *T*_1_ values present in fixed tissue (TR / TI1 / TI2 = 3s / 0.311s / 1.6s; *α*_1_*/α*_2_ = 4°*/*6°);

Finally, for the last sample position, DWI protocol was acquired to provide fiber orientation information (TR / TE = 3.78s / 71.2ms, 256 diffusion-encoding gradient directions, b = 2500 s/mm^2^). Because the formalin fixation process and the reduced temperature of the sample compared to *in vivo* (Room Temperature ≃23°) significantly reduces water diffusivity, the protocol was repeated 20 times to achieve robust fiber orientation information.

### 2.6. Ex vivo data processing

The MP2RAGE contrast is insensitive to transmit and receive B1 fields that vary significantly when rotations as large as 90 degrees were applied to the sample. Therefore, each of the 9 MP2RAGE images from the 9 brain rotations were co-registered to a reference position using FLIRT from FSL [39]. Corresponding transformations were then applied to the ME-GRE data (phase unwrapped using a three-dimensional best path algorithm [40]). Finally, the registered data were normalized following Eq. 9. A DTI was estimated for each DWI and the 20 DTIs were averaged using a log-Euclidean framework [41]. Eventually, the fiber orientation was defined as the main orientation of the average tensor.

A ME-GRE dictionary was simulated for this particular acquisition, and the corresponding deep learning network was trained using the parameter ranges described in Tables 1 and 2.Finally, the microstructure parameter maps (FVF, g-ratio, *χ*_*i*_, 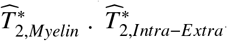, and the relative water weight) were estimated individually for each set of flip angles. This resulted in 6 independent sets of parameter maps, where only the relative water weight term is expected to vary across acquisitions. It was thus possible to compute the mean and standard deviation of the microstructure parameter maps that were expected to remain constant across flip angles to estimate the precision of those measurements.

Finally, the last experiment was performed by using a restricted number of rotations that can be achieved during an *in vivo* experiment. Among the 84 possible combinations of 3 rotations chosen within the original 9 rotations, the 10 that insured the largest fiber orientations ranges were picked. The subsets of *ex vivo* data for the 10 combinations of 3 rotations with a flip angle of 20°, the corresponding dictionaries, and deep learning networks were created, leading to 10 entire sets of brain parameter maps. This was used to compute the mean and standard deviation across different combinations of 3 rotations. Finally, the absolute difference maps between the mean parameter maps with 3 rotations and the original ones with 9 rotations, both with a flip angle of 20°, were estimated.

## 3. Results

### 3.1. Deep learning performance on simulated data

#### 3.1.1. Noise level

Fig. 7(a) shows the dependence of the loss function of the deep learning network for 5 different noise levels as a function of the number of epochs used. After a fast drop during the first 3-5 epochs, the loss function shows a slow decay, reaching a plateau for the noisier signals. Interestingly, the loss functions on the test data (solid lines) have slightly lower values than those on the validation data (dashed line). This difference is attributable to the fact that the validation loss function is averaged along the entire epoch, whereas the test loss function is computed at the end of each epoch. From this analysis we concluded that 20 epochs should be a good compromise between training efficiency and parameter recovery.

**Figure 7:**
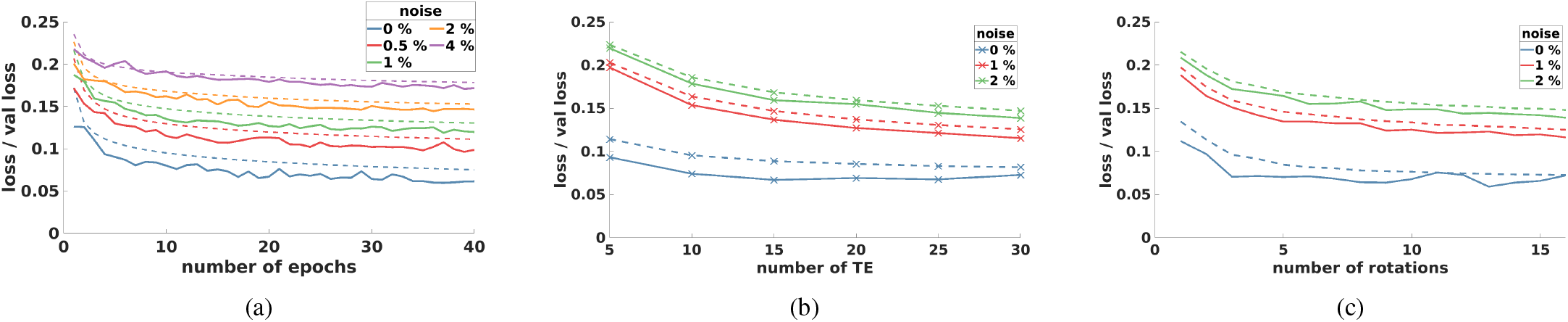
Deep learning training evolution for different noise levels relative to several acquisition parameters. The solid line is the loss function whilst the dashed line is the validation loss function, that represents the same mean absolute error respectively computed on the train and on the test data set. (a) Training along the number of epochs. (b) Training along the number of echoes. (c): Training along the number of rotations.

#### 3.1.2. Echo times

Fig. 7(b) presents the dependence of the loss function on the number of echo times used. It shows that the wider the range of the echo times, the lower the loss function is. The loss function clearly improves between 5 to 15 echoes (corresponding to 49ms), but this improvement becomes smaller once this threshold is passed, even if a plateau has never been totally reached for a signal with noise even after 30 echos. Our simulations did not include any echo time dependent noise, arising from physiological noise or scanner drifts, which are common in gradient echo acquisitions, and would make subsequent echo times less useful for decoding. We postulated that 20 echos would be sufficient for an experimental protocol.

#### 3.1.3. Number of magnetic field orientations

Fig. 7(c) shows that, as expected, the loss functions decrease when the number of rotations for all noise levels increases and it is true for all noise levels. Note that, in the interest of computation time, the subset of rotations might not be optimal for all number of rotations tested (as a subset of the initial 16 orientations was used). Furthermore the specific number/set of rotations depends on the orientation of the fiber of interest. The deep learning benefits from the first 3-6 distinct rotations, similar to what has been demonstrated for susceptibility tensor imaging [42] and for fiber orientation mapping [17], and plateaus after this. In a given acquisition time we can either decide to have an improved SNR per orientation or increased number of rotations. When moving from 1 to 2% SNR levels, this corresponds to a decrease in the acquisition time or number of rotations by a factor 4. Thus, 16 orientations at 2% noise could be acquired in the same time as 4 orientations at 1% noise level. It can therefore be concluded that there is a limited benefit in maximizing the number of orientations beyond 5 as the loss function for 16 rotations at 2% noise was the same as that of 6 orientations at 1% noise. In our acquisitions, we used 10 orientations, to avoid excessive acceleration of each orientation, as this could bring parallel imaging artifacts into play when trying to further reduce the acquisition per orientation.

#### 3.1.4. Selective set of parameters

Fig. 8 shows the performance of the deep learning networks in recovering the various microstructural parameters of what could be considered a typical WM model. Although the average recovered parameters are close to the original ones regardless of the signal noise level, many of the differences would be significant. Particularly, the relative water weight suffers a constant positive bias for all networks and simulated signals. Surprisingly, the standard deviation for all parameters (excluding χ_*i*_and 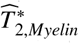) is considerably lower when the deep learning was trained with a 4% noise level rather than the matched noise level. Thus, a dictionary with a high noise level was used in our *ex vivo* experiment presented in the following. When comparing the width of the various distributions, compared to the range used for the training the network (see Table 1), the values of *χ*_*i*_, g-ratio and relative water weight are likely to have the largest biases and noise.

**Figure 8:**
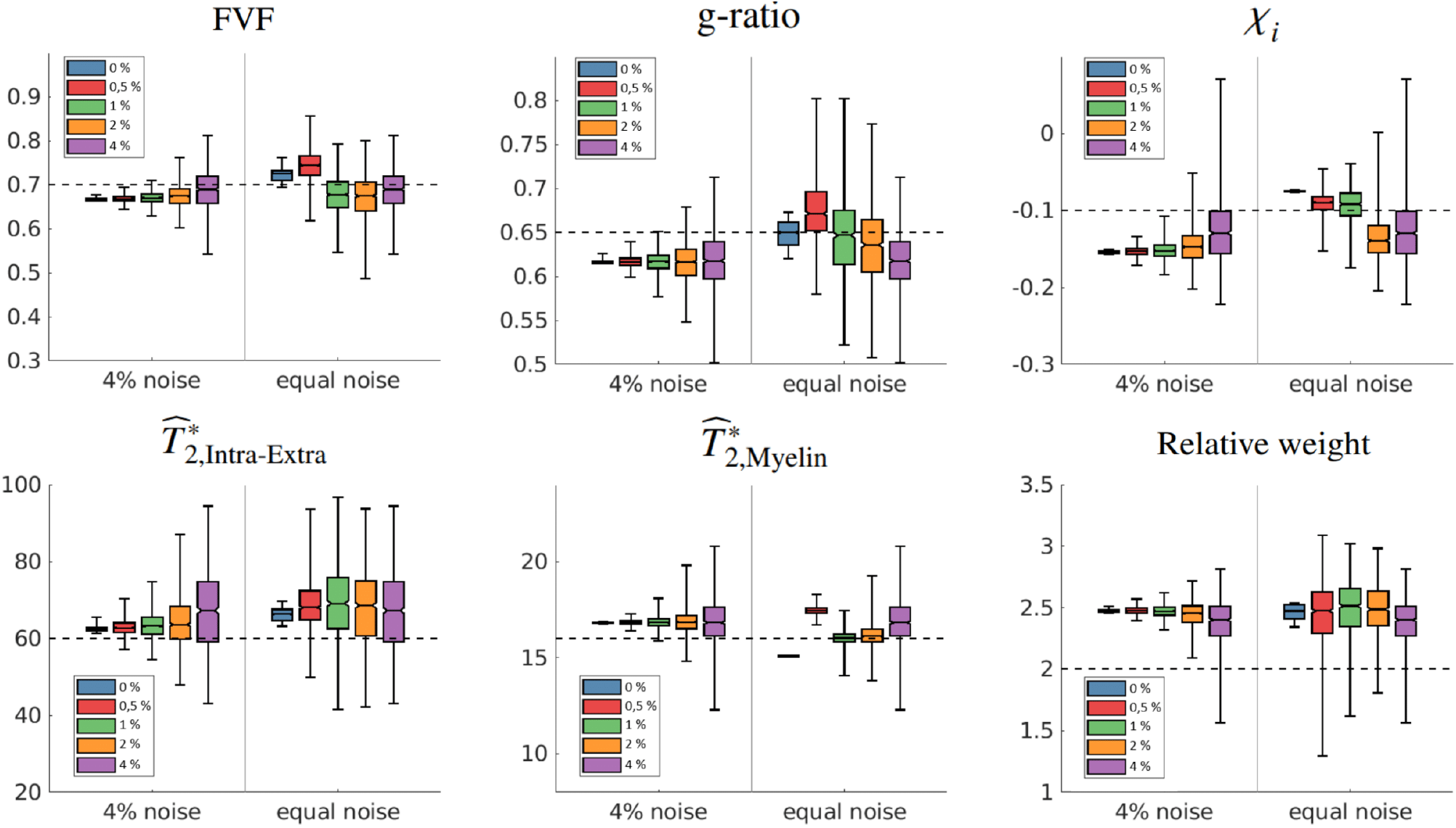
Each box represents the estimation of one parameter recovery for 5 different signal noise levels (0%, 0.5%, 1%, 2%, 4%), the dashed lines represent the correct values. Within a box, the left side use a single deep learning trained with 4% noise regardless of the noise level while the right side use 5 deep learning, each one trained with a noise equal to the signal level.

### 3.2. Ex vivo experiment

Using the deep learning network described on Tables 1 and 2 it was possible to derive 6 microstructural parameters (FVF, g-ratio, *χ*_*i*_, 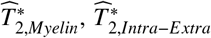, and the relative water weight). For the sake of better visualization, we choose to present the 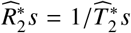 maps instead of the 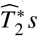 maps. The 6 microstructure parameter maps obtained from the *ex vivo* brain data acquired with a flip angle of 35° estimated with and without the Lorenzian correction are presented in the left and right panels of Fig. 9. WM is clearly discernible from GM and deep gray matter on the FVF, and relative water weight maps. The 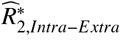 and *χ*_*i*_ maps have weak contrast between GM and WM, while g-ratio is decreased in WM (more myelin surrounding axons and creating dephasing in free water compartment). This observation is particularly interesting because it suggests that, with our modelling, we were able to remove myelin contributions from typically observed 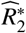 contrast. The sagittal maps show that a higher FVF, lower 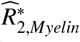 and lower g-ratio in the corpus callosum compared to the rest of the brain.

**Figure 9:**
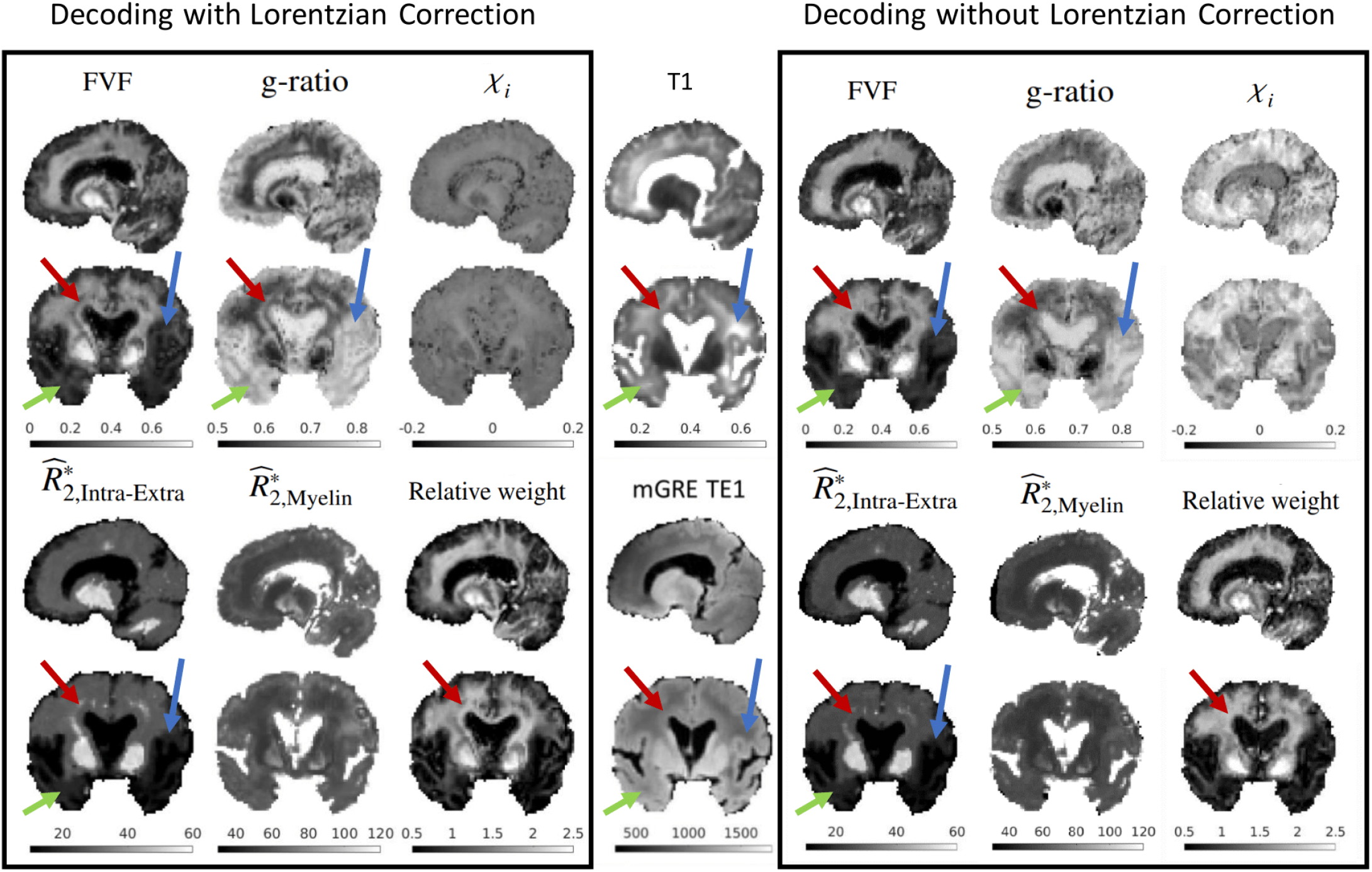
The left and right panels show the 6 parameter maps estimated from the *ex vivo* acquisition with flip angle 35° with the Lorentzian correction and without the Lorentzian correction. The middle column shows a *T*_1_ map estimated from MP2RAGE and downsampled to the resolution of the ME-GRE, and the magnitude of the first echo of a ME-GRE for visual comparison of the contrasts. Each image contrast and parameter map is shown on a sagittal and a coronal slice crossing the globus pallidus. Arrows highlight WM regions where: blue - the microstructural maps correctly reflect tissue properties; green - where the contrast is unexpected given the *T*_1_ maps; red - artifacts are highlighted by using the cylindrical Lorentzian correction in our dictionary.

In WM there are significant variation of contrast in the microstructural maps in the coronal slice. Some follow the same pattern seen on the *T*_1_ maps, very elevated values on the right temporal lobe and above, see blue arrows Fig. 9, that could result from elevated g-ratio and reduced FVF. This suggest its origin to be a fixation artifact or tissue damage that also impacts the observed *T*_1_ values. Note that the long fixation time of the brain sample has resulted in an inversion of the *T*_1_ contrast of white and gray matter (WM having a longer *T*_1_ than GM) with respect to in vivo as well as significant decrease in *T*_1_ values, particularly in deep gray matter, which supports a dramatic tissue changes resulting from the fixation. On the left temporal lobe (see green arrows Fig. 9), this pattern is not reflected in the *T*_1_ maps, but is seen the magnitude image and could either be real or suggestive the breaking down of the decoding process. Red arrow highlights a structure that appears as bright in 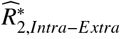 and g-ratio, and dark on FVF maps. This contrast which is thin within most of the slices (as seen on the sagittal cut) is not directly visible on the raw images and is particularly evident when the Lorentzian Correction is used in our signal dictionaries. This suggests that the Lorentzian correction is less appropriate than the standard HCM to characterize this data, it should be noted that this conclusion cannot be extrapolated to fresh tissue and *in vivo* imaging.

Interestingly, CSF presents an almost null FVF along with a high 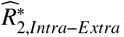, which is to be expected as there are no structures generating an anisotropic signal evolution in this region. The *χ*_*i*_ map estimated with the Lorentzian correction has a very weak contrast with a mean value close to 0 while the map estimated without the Lorentzian correction have mostly positive values within WM with a significant contrast between WM and CSF. Despite the common belief that myelin is diamagnetic, both positive [29] and negative values [17, 13] have actually been reported in the literature. Assuming that phospholipids are diamagnetic, the value of isotropic susceptibility should then be attributed to one or a combination of the following aspects: (i) fixation process that could render the intra and extra axonal spaces more strongly diamagnetic than the myelin sheath; (ii) The not fully understood orientation dependence of the frequency shift of the myelin water compartment. Appendix A presents three different white matter models: the classic HCM, the HCM with a Lorentzian correction, and a layered model. These models have a very different myelin water frequency shift along the B0 orientation that impacts the *χ*_*i*_ estimation (see the maps with and without Lorentzian correction). Thus, the layered model which inverts the angular dependence of the myelin water frequency shift could estimate a positive susceptibility.

One region where the deep learning approach gave unsatisfactory results was in the globus pallidus and the dentate nucleus. These regions are known to be amongst the most iron rich regions in the brain [43, 44], and have therefore very short-apparent transverse relaxation rates in the free water compartment. This was correctly mapped by the 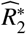 of the intra-axonal compartment, but appears mismapped on the FVF maps. In the latter, the globus pallidus appears as having a large FVF (which is known not to be the case), although the neighbouring putamen appears to be correctly mapped. Surprisingly, the globus pallidus appears as having a similar isotropic magnetic susceptibility to neighbouring WM. It is know that deep gray matter structures are rich in iron, and assuming that this is equally distributed in the intra and extra-axonal spaces, the equivalent field distribution to be generated by our realistic models would require a heavily diamagnetic myelin compartment (beyond our current dictionary limits). Alternatively this result could be explained by such nuclei having a more randomly distributed micro-structure organization that is not well described by our dispersion free tissue models. These observations suggest that further improvement of the realistic model are needed to be able to describe iron rich gray matter.

To demonstrate the robustness of the microstructural parameter map findings with respect to the changes of the acquisition protocol, we analyzed the acquisitions with different flip angles separately. The microstructural parameters should not depend on the flip angle, except the relative water weight that is linked not only to the proton density but also the relative saturation of each compartment (which depends on *T*_1_, TR and the flip angle). The mean parameter maps, estimated with the Lorentzian correction, showed overall the same characteristics than the ones presented previously for flip angle 35. In supplemental material it can be seen that the standard deviation of the microstructure parameters FVF, g-ratio, *χ*_*i*_, intra-axonal, 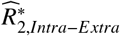 and 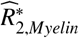 are small when compared to the decoded maps, the only exception being the relative weight map.

Fig. 10 demonstrates the possibility of decoding the microstructure parameter maps, estimated with the Lorentzian correction, from only 3 brain orientations. The mean parameter maps highlight the expected brain structures, and are in good agreement with our ground truth (obtained from 9 brain orientations). The standard deviation maps estimated across 10 combinations of 3 different rotations (middle row), reveal very low values when compared to the recovered values (note that the colorbars of the standard deviation maps are significantly reduced with respect to those used to show the decoded parameters). Thus, the process is robust to the specific set of orientations used. Interestingly, the contrast seems lower compared to the parameter maps obtained with 9 rotations, in particular within deep gray matter, as illustrated by the absolute difference maps (see bottom row). As mentioned earlier, this is the region where our model is failing to describe the microstructure properly.

**Figure 10:**
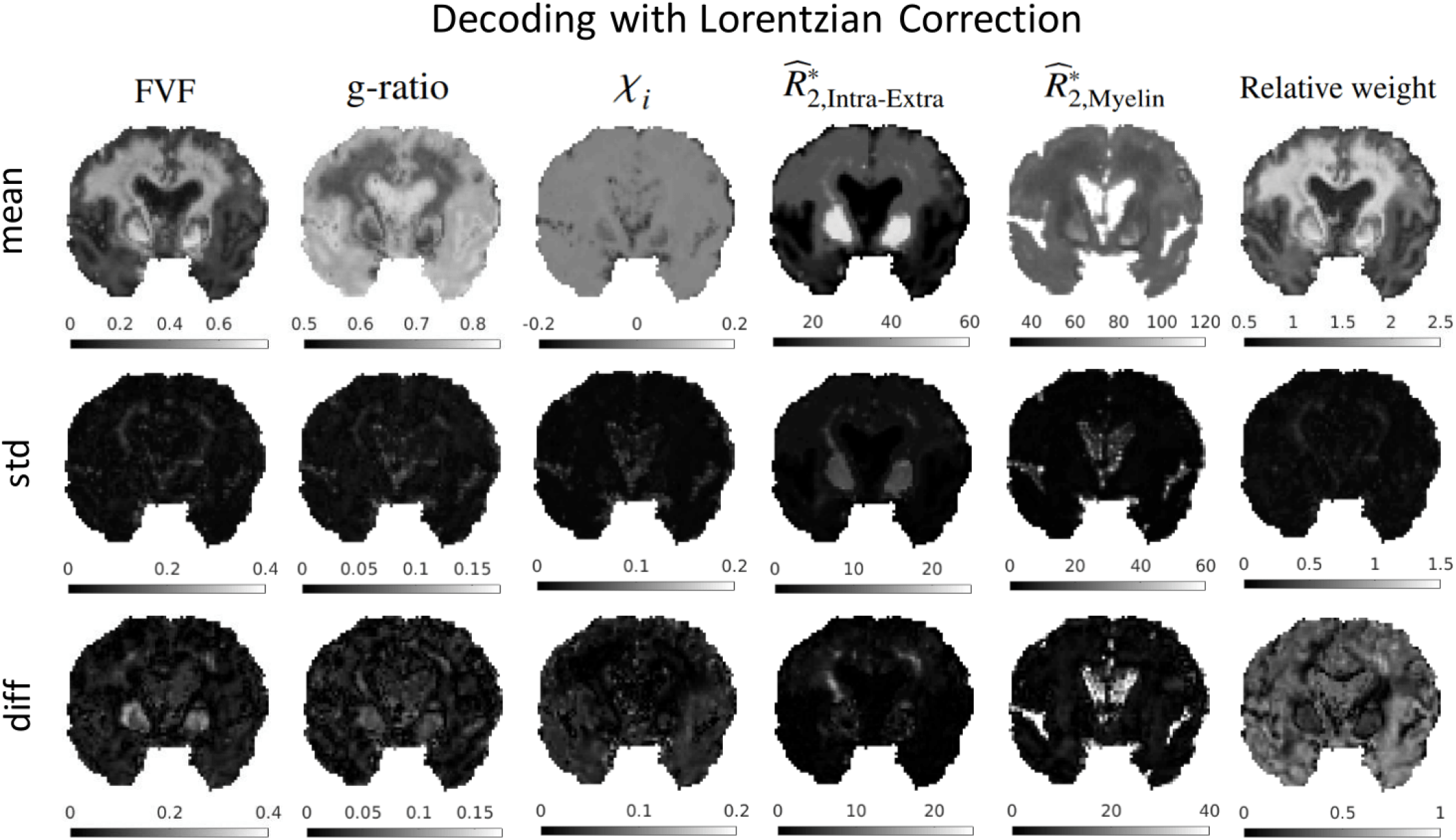
Coronal slices of the 6 decoded microstructure parameter maps (FVF, g-ratio, *χ*_*i*_, 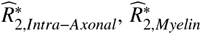, and relative weight shown from 1st to 6th column) obtained from the ME-GRE with flip angle 35°. The top row shows the averaged parameter maps obtained using 10 distinct dictionaries trained with only 3 orientations. The middle row shows the standard deviation across the various decoded maps obtained from the different subset of 3 orientations. The bottom row shows the difference to the absolute difference of the obtained maps in respect to the our “ground truth” (maps decoded from data containing 9 different head orientations).

## 4. Discussion

### 4.1. White matter models: promise and limitations

We introduced a pipeline to create a simple yet realistic biophysical model to simulate the MRI ME-GRE signal. These WM models contain real axon shapes and a g-ratio variability similar to what is reported in tissue samples, and have varying levels of FVF within themselves as a result of the axon packing approach. With the realistic WM models available for microstructural quantification, it can be used as an alternative means in contrast to the analytical expression of WM microstructure in parameter mapping which can lead to measurement bias as previously reported [14]. Yet, some effects are deliberately overlooked: (1) diffusion within the compartments, (2) chemical exchange, (3) compartmentalization of water within the myelin sheath and (4) other sources of susceptibility perturbations beyond the myelin sheath.

Diffusion has been demonstrated to have a minor effect in WM models based on EM data [14] when compared to the hollow cylinder model or simple cylindrical perturbers [45]. Chemical exchange between myelin water and myelin protons results in frequency shift, and thus, can be accounted for by adding an exchange term in the HCM [23]. The size of this frequency offset term has been reported to be of 0.02 ppm in the corpus callosum [17], but models have been proposed that would make this offset depends on the number of myelin layers and therefore varies throughout the brain and fibre bundles [46]. Yet, chemical exchange has been demonstrated to have a larger impact when measuring the longitudinal relaxation in WM (a process that is much slower than time scales explored here).

This is the main reason why longitudinal relaxation mechanisms were up to now hidden in the weight parameter.

In Appendix A, we compared the classical HCM [17] implicitly used in previous WM models [14], with a layered model where the source of susceptibility (phospholipids) is spatially separated from the source of signal [15, 30] and the new implementation of the Lorentzian correction (see Eq. 6) used in our realistic WM models. The main aspects addressed in this comparison were the frequency distribution of the different compartments. As has been analytically described the field perturbations are equivalent in the intra-axonal and extra-axonal compartments. Conversely, strong differences exist in the myelin water compartment for the three models, which predict opposed angular dependence of the mean frequency shift of this compartment as a function of the axon orientation with respect to the magnetic field. The results described in Fig. A.12 of the Appendix A suggest that the cylindrical Lorentzian correction within the myelin compartment would better fit experimental data without requiring an exchange mechanism [17] or the hop in hop out of water across myelin layers described in [15]. Introducing the new correction in our network resulted in an isotropic susceptibility that was more diamagnetic than otherwise (see Figure 9) but seemed to enhance some decoding artifacts in some of the resulting micro-structural maps.

The extra-axonal compartment currently includes everything that is found outside the axon. More classes with specific properties could be used, particularly: free water (CSF and interstitial spaces); blood vessels; bound-water compartment (that represents the water bound to macromolecules present in cell walls and organelles [47]), and iron accumulated in ferritin, amongst other. Blood vessels represent a very small fraction of the tissue volume (1-4% in WM and GM, but venous blood, which is deoxygenated, has a much larger susceptibility different to free water than myelin) and tends to follow the orientation of WM axon bundles [21]. This is expected to introduce some degree of 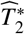 anisotropy that would act as a confound in our *ex vivo* experiment. Ferritin, which is known to be strongly paramagnetic, can be found everywhere in the brain (with increasing quantities found from WM, GM to deep gray matter where it can be found in large quantities [48]). On our current implementation, iron is expected to be equally distributed in the intra- and extra space. As a result, ferritin will be mapped as a reduction of the 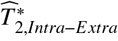 and the isotropic magnetic susceptibility attributed to the myelin compartment is effectively the difference between the susceptibility of the myelin and the free water compartments where there might be ferritin inclusions. Note that in the case of high ferritin concentration our current cylindrical Lorentzian correction will be overestimated.

### 4.2. Dictionary and deep learning

Many of the simplifications used in our WM models arise from the need to restrict the number of parameters associated with our network. The size of a dictionary, which in this study had 7 dimensions (see Table 1), is around 10 GB. Moreover, an increase in the number of variables mapped by the network will result in an increased noise level of the parameters estimated. We believe we have restricted the models to the most relevant parameters. In particular, we have considered FVF and g-ratio inherent to the model, as described previously the extra-axonal space can have various types of constituents, thus the extra-axonal 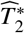 cannot be fixed. We choose to free *χ*_*i*_ (allowing this to incorporate magnetic susceptibility in the intra-/extra-axonal compartment) and to fix *χ*_*a*_, as the major contribution to the magnetic field perturbation comes from the isotropic susceptibility [49]. The compartment water weights were represented by a single variable, the relative water weight that includes the water proton density as well as the degree of *T*_1_-weighting (and chemical exchange) of each compartment. If the myelin sheath is considered having the same properties all over the brain, that allows to fix the 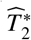 of myelin and release the anisotropic susceptibility *χ*_*a*_ which was reported to be ranging from − 0.15 to − 0.09 ppm [18]. A potential direction for future work is to investigate different sets of parameters. For example, the myelin water concentration may be linked to the susceptibility of the myelin sheath, bearing in mind that the magnetic susceptibility of the phospholipids and water are both known. This would benefit from some of the insights gained from our Appendix A.

Our deep learning network is robust and systematically converges for each dictionary associated to an experiment with multiple orientations, as illustrated in Fig. 7. However, extensive manual fine-tuning of the network hyper-parameters was required to achieve this level of agreement. A more systematic approach, while potentially desirable, would need an excessively long computation time. The *in silico* analysis (see Fig. 8) shows that a dictionary trained with a higher noise level is more robust to noise amplification than a dictionary with matched noise levels. This was attributed to the noise allowing to smear our differences associated with the fact that our “realistic model” produce different signals (see Fig. 5) and none of them really corresponds to the actual WM mapped. An interesting experiment would be to assess the performance of a dictionary including all different noise levels, mimicking closer to the signal found in the brain, where regions further away from the receiver coils are bound to have a lower SNR. It was observed that the level of noise remains within the range that differentiates our 2D models from a real 3D WM for a relatively wide range of dispersion values, which effectively makes our network more generalizable. Nevertheless, once the neural network is trained, it can provide much faster processing speed when compared to transitional voxel-wise data fitting approach (few seconds vs several minutes with typical 2-mm isotropic whole-brain coverage data for gradient echo MWI [50].

### 4.3. Ex vivo experiment

The human brain scanned on our *ex vivo* experiment was fixed in formalin for 7 months prior to the experiment. It is well known that the microstructural tissue properties change throughout the fixation process, and the final properties of the tissue depend on: the *post-mortem* fixation delay, the fixation time, the concentration of formalin and the temperature history [51, 52, 53]. The *T*_1_ map presented in Fig. 9 shows particularly small values revealing strongly fixed tissues where water has a reduced mobility. The mean ADC in WM found was 0.3 *mm*^2^ *s*^−1^ while a normal *in vivo* value would be above 0.8 *mm*^2^ *s*^−1^ [54]. While it had already been demonstrated that adding diffusion to realistic models of WM was not relevant when trying to model the GRE signal [14], for fixed tissues this should be even more so. One risk of using fixed tissues is that protein binding might change the sizes of the different compartments and tightness of the myelin packing, making our realistic models less valid. Fresh tissues do not present such problems (reduced diffusion and fixation artifacts) and could be an alternative option. However, our current protocol took 8h, without the DWI. In such a time window, fresh tissues would not be sufficiently stable to assume constant microstructural properties over time [55]. Thus, it was necessary to use fixed human brain for this proof of concept microstructural parameter decoding experiment, although the fixation time could be reduced to 6-10 weeks would make our findings more comparable to what is found *in vivo*.

## 5. Future work

### 5.1. Ground truth validation

We have demonstrated the feasibility of incorporating realistic models to measure WM microstructural properties. While the recovery of the microstructural parameters generated follows the general expectations for FVF, 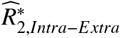, g-ratio, it would be important to validate them by an independent method is future experiments. One possible avenue is to perform histology on excised samples after the MRI experiment which could provide ground truth of microstructural properties. Amongst the potential candidates: CLARITY is an optical 3D imaging method combined with a tissue clearing technique which can provide neuro density, fibre orientation distribution and cell types [56]; X-ray microscopy can also be used to generate an entire 3D view in a non-destructive way [57]; 3D transmission electron microscopy, as used to generate the axon models, can provide excellent resolution to quantify myelin volume and axonal orientations. However, using histology as a means for method validation has to be interpreted carefully as the tissue preparation for processes can change the intra and extra-axonal water content and relative volume. A direct comparison of the obtained microstructural parameters obtained *ex vivo* on fixed tissue with *in vivo* acquisitions should be avoided. In a preliminary work (data not shown), we replicated the fixation formalin process and the MRI experiment with a porcine brain sample, from which small WM samples were excised for 3D EM analysis. We observed significant degradation of the myelin sheath for a number of axons where the myelin sheath appeared unpacked. Such tissue change could result in a decrease of g-ratio, *χ*_*i*_, as well as a decrease of myelin 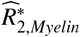 and proton density with respect to *in vivo* imaging. Yet, given the lengthy MR acquisition performed, using a fresh tissue sample would not be feasible.

In this study, we did not compare our proposed method to any conventional imaging methods for WM microstructural quantification. Our realistic WM model driven with deep neural network provides a set of microstructural properties that is unique making direct comparison to other microstructural quantification methods not straightforward. For example, myelin water imaging [9] might be a combination of FVF, g-ratio and the myelin weight term derived by our method, the NODDI obtained intra-axonal volume fraction would be a combination of FVF and g-ratio. Additionally, the structural alteration of *ex vivo* samples hinder the robust applicability of the conventional methods in *ex vivo* imaging: previous studies have shown that the 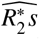 of myelin and intra-/extra-axonal water become less distinguishable in *ex vivo* data [58] and, as in our data, the reduced water diffusivity in *ex vivo* samples makes the extraction of information beyond main fiber orientation extremely challenging. Future work will address such comparisons *in vivo*, where our deep learning methods could benefit from additional diffusion modelling information.

### 5.2. Application

The current implementation of the network (based on ME-GRE data acquired with one single flip angle and information regarding the main fiber orientation) requires at least 3 head orientations with respect to the main magnetic field. Although this limits its applicability *in vivo* it is comparable to the requirements of other magnetic susceptibility related methods such as COSMOS [59] and Susceptibility Tensor Imaging [60] and compares favorably to magnitude and frequency-based fiber orientation estimation [17]. Two possible solutions recently introduced to raise this degeneracy in the context of myelin water imaging is to explore the difference between the *T*_1_ of myelin and free water by using various GRE acquisitions with different *T*_1_- weighting and use additional information from DWI regarding the relative size of intra and extra-axonal components [61].

The approach presented in this work may find applications in the imaging of myelin water with gradient-echo-based acquisitions [62, 10]. Traditionally, myelin water imaging in gradient-echo-based experiments tries to fit 9 independent parameters: three independent signals (separate amplitude, decay rate and frequency shift) for each of the three compartments (intra- and extra-axonal water and myelin) to a ME-GRE signal. The main shortcomings of this approach are that: the model is known to be simplistic (even the simple HCM predicts more complex signal evolution than 3 overlapping exponential signal decays) [23] and the fitting procedure is ill-conditioned. In the work presented here, we have shown with simulations and data that we can obtain acceptable results with as few as 3 orientations with the advantage that we obtained the most relevant microstructural information. One of the common findings in myelin water imaging is, as in our *ex vivo* results (see Figs. 9, 10) an overestimation of the myelin water compartment in deep gray matter [62, 63], where the limitations of the tissue model become evident. One approach is to use advanced diffusion modelling priors that, for example, quantify intra- and extra-axonal water fractions [64, 65] or describe each voxel as being an overlap of various fiber orientations [66]. Both these approaches have been successfully been demonstrated recently in the context of *in vivo* myelin water imaging [61], but because of the poor quality of the *ex vivo* diffusion data could not be pursued. Finally, addressing specifically the erroneous fitting in deep gray matter, it is foreseeable to integrate this methodology with QSM [59], in which case the additional information on regions that are high in iron load could avoid applying a model that does not describe this regions appropriately.

## 6. Conclusion

In this paper, we developed an open source toolbox ^2^ to generate 2D WM models with controlled microstructural properties such as fiber density and variability in the g-ratio using publicly available electron microscopy data. Such models are used to estimate the corresponding field perturbation and derive the ME-GRE signals. Although our WM models are limited to 2D, we have demonstrated that they can be satisfactorily used to simulate 3D structures with a relatively high range of dispersion. Finally, dictionaries of ME-GRE signals for 6 different parameters (FVF, g-ratio, *χ*_*i*_, 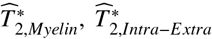, and the relative water weight) associated with WM properties at a sub-voxel level were created. This single acquisition dictionaries can then be combined depending on the multiple rotation strategy used in the experimental protocol to create a better conditioned decoding problem and train a deep learning network able to decode microstructural parameters. We performed several tests to assess the quality of the sub-voxel parameter recovery using our network, depending on the number of sample rotations, echo times used and noise added to the library. Unsurprisingly we found that the network performs better as more data are given as input. Thus the number of rotations and echo times should be maximised in a given acquisition window. However, because of the large variations between different WM models used in the training process, it is advantageous to train the network with a level of noise higher than that of the available data. The network was demonstrated through an *ex vivo* experiment performed using gradient echo data acquired at multiple brain orientations with respect to the main magnetic. We were able to obtain promising FVF, g-ratio, 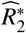 maps that showed the expected variations through out most brain structures such as the CSF, GM, WM and the corpus callosum. The parameter values (except for *χ*_*i*_) follow the expected patterns and were robust for different acquisition protocols and reduced number of brain orientations.

## Supporting information

Supplemental Video: White matter model creation

## 7. Aknowledgements

The authors would like to thank Professor Karla Miller and Dr. Michiel Kleinnijenhuis for providing us with the 3D electron microscopy data and its segmentation that was used in the Appendix B. This research was supported by the Nederlandse Organisatie voor Wetenschappelijk Onderzoek (NWO), Grant/Award FOM-N-31/16PR1056 that sponsored the positions of Renaud Hédouin and Kwok-shing Chan. The authors would also like to acknowledge the fruitful discussions on the topic of this research with Prof. David Norris and the insights brought by the reviewers regarding the compartmentalization of water in the myelin water compartment.

## Appendix A. Impact of compartmentalization of water within the myelin sheath

### Appendix A.1. Presentation of the model

In this appendix, we compare three distinct infinite cylinder white matter models to address the question of how to better model the compartmentalization of water in the myelin compartment: - the classic HCM [17], which considers the myelin sheath with water as one single compartment; - the HCM in which a cylindrical Lorentzian correction [29] was applied to the myelin water compartment as described in Eq. 6; - a layered model that divides the myelin sheath in several phospholipid layers (that are the source of susceptibility) interleaved with 10 myelin water layers (source of the signal) [15, 30];

To describe the layered axon model, the myelin compartment was replaced by phospholipid and water compartments. To ensure that the model does not alter the intra and extra-axonal fields generated by the standard model, the volumetric isotropic and anisotropic susceptibility are set to −0.1*/w*_*P*_, where *w*_*P*_ is the volume fraction of the phospholipid layers in the myelin compartment (1 − *w*_*MW*_, where *w*_*MW*_ is the myelin water fraction). Finally the 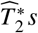 of the phospolipid and myelin water were set to 0.5 and 15 ms. The cylinder models were simulated in a 2000 × 2000 grid, to avoid numerical errors in the presence of large number of layers.

Following the process described in Section 2.2, the field perturbation and the corresponding ME-GRE signal were simulated using the 3 models (with the same FVF and g-ratio) while two different myelin water fraction (in the standard HCM model *weight* matched the layered water volume fraction).

### Appendix A.2. Field perturbations in the layered WM model

The field perturbation and their corresponding frequency distributions of the 3 models are shown in Fig. A.11. As analytically predicted [15], the intra-axonal and extra-axonal frequencies have similar distribution for the 3 models, but the myelin water has very different frequency distributions. Because in the models shown the susceptibility of the axon has been matched, the field within the phospholipid layers is increased, while the water compartment shows the opposite behaviour. The cylindrical Lorentzian correction to the myelin compartment of the HCM, see dashed lines in the histogram, has a much milder effect when the axon is perpendicular to the magnetic field (small reduction of the frequency shift), and has the same effect as the layered model when axon is parallel to the magnetic field (in which case the water is on resonance).

To better understand the relationship between the myelin water frequency in the three models Fig. A.12 a and b show the mean frequency of the myelin water compartments as a function of the orientation with respect to the magnetic field, *θ*. As predicted analytically, the HCM and the layered model have opposed frequency shift dependence on *θ*. As noted earlier, the layered model and the Lorentzian cylinder corrected version of the HCM do not show a frequency shit of the myelin water compartment when the axon runs parallel to the main magnetic field. The impact of changing the myelin water volume fraction (see Fig. A.12 a and b) does not play a significant role. But it should be noted that, when increasing the water volume fraction, the magnetic susceptibility of the phospholipid compartment was increased in the layered model, which is not the case in reality where the susceptibility is a property of the phospholipid bilayer.

It is interesting to note that the HCM with the Lorentzian correction in the myelin water compartment has the experimentally found behaviour where the myelin water has no frequency shift when WM is running parallel to *B*_0_ and a positive frequency shift when perpendicular to *B*_0_. This is achieved without the need to consider hop in and and hop out mechanisms as is the case for the layered model [15]. Because of its simplicity and straightforward application to our realistic WM models, this was the model used in our study.

**Figure A.11:**
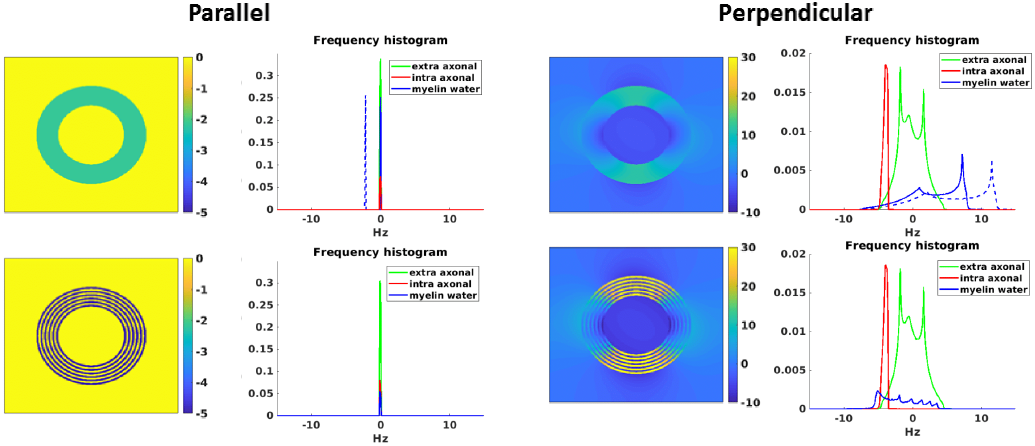
Field perturbations and frequency distributions of the HCM (top row), and the Layered model (bottom row), for a parallel (left) and a perpendicular (right) orientation of the axon relative to the main magnetic field orientations. On the top row, the blue dashed line represents the myelin histogram without the Lorentzian correction while the blue solid line represents the myelin histogram with the Lorentzian correction. The frequency histograms only show the frequency distribution in the compartments with water signal. The reduction in area of the myelin water distribution reflects its decrease in volume fraction.

## Appendix B. 3D WM model

### Appendix B.1. Presentation of the 3D WM model

With a view to validating the ability of the 2D realistic models developed to describe the 3D structures encountered in a WM voxel, we compared the signal associated to the 2D models to those of a real 3D WM sample. A segmented [67] 3D EM of a genu of a sagittal mouse corpus callosum Section was used for this comparison. The resolution of the initial 3D EM dataset was of 7.3×7.3×50 nm, which was subsequently down sampled by a factor of 7 resulting in a quasi isotropic resolution 51×51×50 nm. The FOV of the segmented piece was 20×20×20 *µ*m (represented on a matrix of 400×400×400). Using the segmentation 3D EM data, the FVF and g-ratio were computed to be 0.51 and 0.67 respectively. Additionally, because the 3D model does not consist of infinitely long structures that are parallel, the fiber dispersion was computed with respect to the average fiber orientation [68], and found to be low *σ* = 0.04. In addition to this original model, to study the impact of higher dispersion, 60 axons within the 3D model were selected to create a fiber orientation dispersion of *σ* = 0.4. A mask surrounding the selected axons was used to ensure microstructural parameters remained equivalent to those of the whole sample (FVF = 0.51 and g-ratio = 0.67). The 3D signal was computed only within the mask and selected axons.

**Figure A.12:**
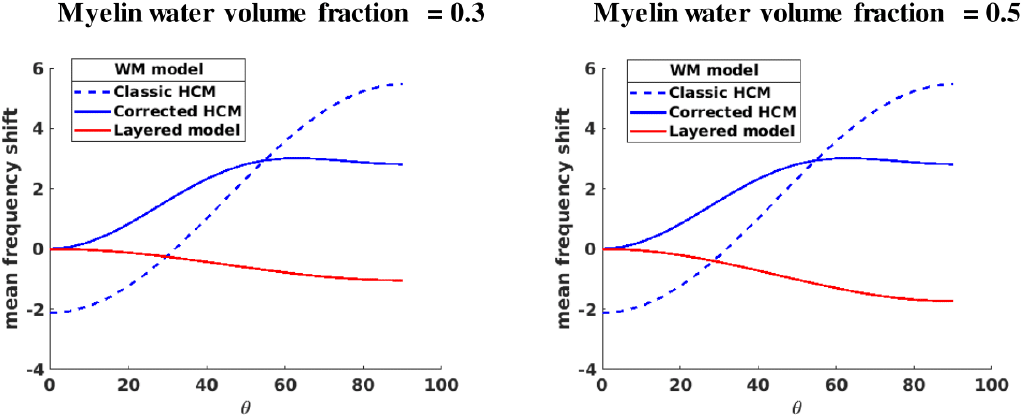
Plots of the mean frequency shift of the myelin water compartment as a function of the orientation with respect to *B*_0_ for the HCM without the cylindrical Lorentzian correction, the HCM with the cylindrical Lorentzian correction and the layered model.

**Figure B.13:**
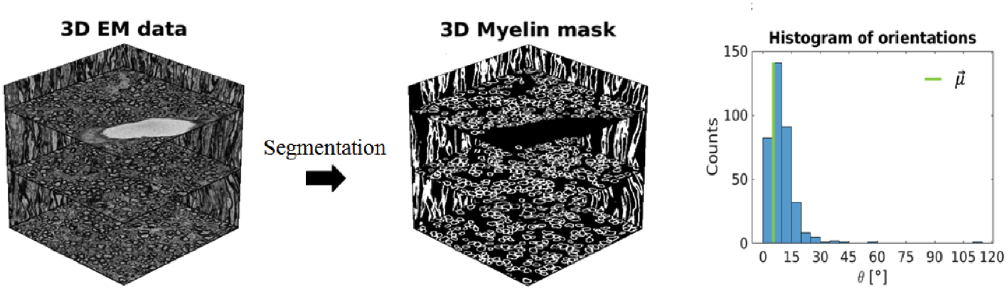
Raw 3D EM data and myelin segmentation of size 400×400×400. Frequency histogram of the computed axon orientations present in the EM model and the average orientation, 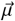

The magnetic susceptibility tensor, *X*_*R*_, was calculated with respect to the orientation of the phospholipids inside the myelin sheath, using a 3D variant of the process described in the methods section. The tensor map obtained was then used to calculate the magnetic field perturbations in 3D, Δ*B*_0_(*X*(*r*)), as described in [28]. These processes are straightforward extensions of the 2D case and their implementation is available in our toolbox.

### Appendix B.2. Comparison between 2D and 3D field perturbations

To simulate the fiber dispersion within the 3D samples, an artificial dispersion was introduced into the 2D models by computing the field perturbation for 100 different main magnetic field orientations according to the von-Mises-Fisher distribution [69]. The final signal is the sum of signal from 2D models with the 100 different orientations with respect to the main magnetic field.

The 3D models were compared to 10 realistic 2D models, created as described in the methods section, using similar microstructural parameters to those of the 3D samples. Four different dispersion values (*σ* = 0, 0.2, 0.4, 0.6) were simulated. The ME-GRE signals were computed for both 2D and 3D models, with the parameter used in Fig. 1 for TE = 1:1:80 ms. Finally, the 2D and 3D signals were normalized and compared using the root-mean-square-error (RMSE) computed according

to:

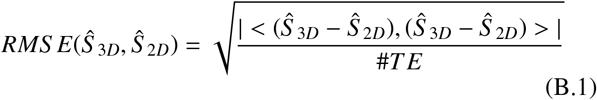

where <.,.> is the complex dot product, |.| the absolute value operator and #*TE* is the number of echos.

Fig. B.14 shows the signal RMSE between the 2D and 3D models as a function of the orientation of the main magnetic field. In each plot, various 2D simulated signals with different dispersion levels are compared to (a) the origin al 3D model (b) the 3D model with high dispersion. The 2D models with lower dispersion (0, 0.2) consistently match that 3D signal with RMSEs below the 2.5%, which is small when taking into account the 4% noise added to the training of our deep learning network used in our *in silico* and *ex vivo* experiments (see Section 2.4.1 and Section 2.6). For the high dispersion 3D model (Fig. B.14b), the 2D models with high dispersion (0.4 and 0.6) have the lowest RMSE for all magnetic field orientations. When no dispersion is used in the 2D models, the RMSE stays below 5%. The two 3D models considered are best represented with 2D models with similar or slightly higher dispersion values. This finding could be attributed to the additional dispersion associated with each axon that changes direction throughout the 3D model and that is not taken into account in the current dispersion computation.

**Figure B.14:**
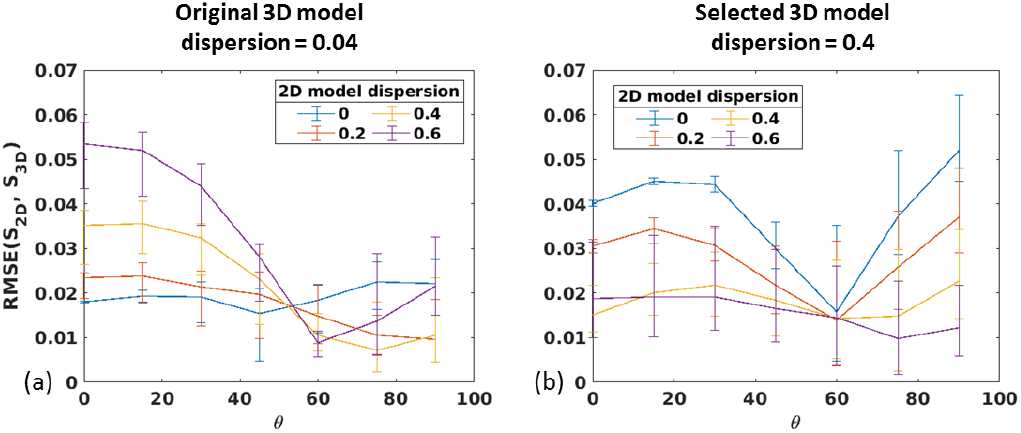
Plots of the RMSE between the Signal of the 2D models using 4 different dispersion levels and Signal of the 3D models as a function of the orientation of the main magnetic field. In a) the original 3D model with low dispersion (0.04) and in b) the 3D model with high dispersion (0.4), which is used as ground truth. The error bars represent the standard deviation across 10 different realistic 2D WM models created

To conclude, the 2D models developed based on separate library of axons accurately represent a real 3D WM model. In the future, it could be considered to add dispersion to the 2D models to better represent a WM region with higher dispersion values that could be measured independently with DWI. In *ex vivo* acquisitions, the quality of DTI data is severely hampered (reduced diffusion constant and reduced 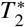), and from our data it was not possible to apply more advanced diffusion models that can decode this quantity. However, even without dispersion, the RMSE consistently remained under 5% while 4% noise is added to our dictionary when training the deep learning network, which suggests that this might not have a large impact.

A situation not considered here and that should have a larger impact are crossing fibers. Fiber dispersion, discussed above, accounts for the spread of the fiber orientations within a bundle of axons while the fiber crossing represents two or more bundles of axons. Significant work on the diffusion community has been devoted to this topic [70]. This could be studied as future work assuming that such a 3D EM dataset exists.

## Footnotes

1 https://osf.io/sgbm8/

2 https://github.com/rhedouin/Whist

